# Evaluating metagenomic assembly approaches for biome-specific gene catalogues

**DOI:** 10.1101/2021.09.30.462683

**Authors:** Luis Fernando Delgado, Anders F. Andersson

## Abstract

For many environments, biome-specific microbial gene catalogues are being recovered using shotgun metagenomics followed by assembly and gene-calling on the assembled contigs. The assembly can be conducted either by individually assembling each sample or by co-assembling reads from all the samples. The co-assembly approach can potentially recover genes that display too low abundance to be assembled from individual samples. On the other hand, combining samples increases the risk of mixing data from closely related strains, which can hamper the assembly process. In this respect, assembly on individual samples followed by clustering of (near) identical genes is likely preferable. Thus, both approaches have pros and cons and it remains to be evaluated which assembly strategy is most effective. Here, we have evaluated three assembly strategies for generating gene catalogues from metagenomes using a dataset of 124 samples from the Baltic Sea: 1) assembly on individual samples followed by clustering of the resulting genes, 2) co-assembly on all samples, and 3) mix-assembly, combining individual and co-assembly. The mix-assembly approach resulted in a more extensive non-redundant gene set than the other approaches, and with more genes predicted to be complete and that could be functionally annotated. The mix-assembly consists of 67 million genes (Baltic Sea gene set; BAGS) that have been functionally and taxonomically annotated. The majority of the BAGS genes are dissimilar (<95% amino acid identity) to the Tara Oceans gene dataset, and hence BAGS represents a valuable resource for brackish water research.

**IMPORTANCE:** Several ecosystem types, such as soils and oceans, are studied through metagenomics. It allows the analysis of genetic material of the microbes within a sample without the need for cultivation. When performing the DNA sequencing with an instrument that generates short sequence reads, these reads need to be assembled in order to obtain more complete gene sequences. In this paper, we have evaluated three strategies for assembling metagenome sequences using a large metagenomic dataset from the Baltic Sea. The method that we call mix-assembly generated the greatest number of non-redundant genes and the largest fraction of genes that were predicted to be complete. The resulting gene catalogue will serve as an important resource for brackish water research. We believe this method to be efficient also for generating gene catalogs for other biomes.

## INTRODUCTION

High-throughput sequencing has led to the establishment of the metagenomic field, allowing the direct analysis of genetic material contained within an environmental sample (1). This approach offers a detailed characterization of complex microbial communities without the need for cultivation. It can be used to address questions like *which* microorganisms are present, *what* are they capable of doing, and *how* do they interact. Metagenomics has been used for studying several ecosystem types, such as soils, human gut and oceans (2–4)

For many environments, biome-specific gene catalogues have been recovered using shotgun metagenomics, followed by assembly and gene calling on the assembled contigs. Examples are the Integrated Reference Catalog of the Human Microbiome (4) and the Tara Oceans gene catalog (2). Gene catalogs facilitate the discovery of novel gene functions and gene variants. Annotated gene catalogs can also serve as genomic backbones onto which sequencing reads from metagenomes and metatranscriptomes, as well as mass-spectrometry spectra from metaproteomics, can be mapped, which enables fast and accurate taxonomic and functional profiling with such datasets.

The assembly can be carried out either by co-assembling reads from all the samples (or groups of samples) or individually assembling reads from each sample. The co-assembly approach has the advantage that some genes displaying too low abundance to be assembled from individual samples may reach enough coverage to be recovered. However, combining data from many samples often means mixing data from a diversity of closely related strains (from the same species). This fine-scale genomic variation can compromise the assembly process because the de-Bruijn graph will include many alternative paths. Consequently, the assembler may decide to break the graph in smaller pieces, which can result in fragmented genes.

An alternative approach is to perform assembly on each sample individually. The individually assembled samples approach will minimize the mixing of data from different strains and therefore potentially result in more completely assembled genes, at least for fairly abundant genomes. However, another problem arises, which is that (more or less) identical genes from multiple samples will be reconstructed. To serve as a reference dataset, it is desirable to have a non-redundant set of genes. Sequence redundancy removal can be achieved by clustering the gene sequences (or their protein translations (5)) resulting from the different assemblies based on sequence similarity, using some cut-off criteria. For each gene cluster, a representative sequence is then chosen based on e.g., gene completeness, centrality in the cluster, or abundance in the dataset.

Recently, a Baltic Sea specific gene catalog with 6.8 million genes was constructed based on the metagenomic data from 81 water samples spanning the spatiotemporal gradients of the Baltic Sea (6). For the construction of the Baltic Sea specific gene catalog, all the 2.6 billion (i.e., 10^9^) reads were co-assembled and genes called on all contigs >1,000 bp. While this gene catalogue has established itself as a useful resource for analysing metagenome and metatranscriptome datasets from brackish environments (7–11), only ca 10% of the shotgun reads from a typical Baltic Sea metagenome sample are mapping to genes with a functional annotation (6). A reason for the seemingly low coverage could be that the co-assembly approach has resulted in a fragmented assembly. A more comprehensive reference gene catalogue would hence be desirable for this environment. In this study, we conduct an extensive comparison of three assembly approaches on an expanded set of metagenome samples from the Baltic Sea, and present an updated gene catalogue for the Baltic Sea microbiome.

## MATERIALS AND METHODS

### Metagenome samples

Five previously published sample sets (6, 7, 12) were used in this study. The sampling locations are shown in Fig. S1 and a brief description of sample retrieval and sequencing is given in Table S1; for further details we refer to the original publications. Sequencing of all sample sets was conducted using Illumina Hiseq 2500.

### Pre-processing of reads

Removal of low-quality bases was performed earlier (7) using Cutadapt (13) (parameters -q 15,15) followed by adapter removal (parameters -n 3 –minimum-length 31). The resulting read files were thereafter screened for PCR duplicates using FastUniq (14) with default parameters.

### Assembly

Individual assemblies on the 124 samples were performed earlier (7), using MEGAHIT (15) v.1.1.2 with the “--presets meta-sensitive” option. For the co-assembly conducted here, all pre-processed reads were first combined and normalised using BBnorm of BBmap v.38.08 (https://sourceforge.net/projects/bbmap/) with the following parameters: target=70, mindepth=2, prefilter=t. Also, the normalized read set was too extensive to allow co-assembly with the tag “presets –meta-sensitive” with MEGAHIT. Therefore, they were assembled with “--presets meta-large” (using MEGAHIT v.1.1.2), as recommended for complex metagenomes in the MEGAHIT documentation.

### Gene prediction

Genes were predicted on contigs (from the co-assembly and from the individual assemblies) using Prodigal (16) v.2.6.3 with the -p meta option.

### Protein clustering

Clustering of the proteins stemming from the different samples for the individual-assembly, and from the co-assembly for the mix-assembly strategy, was performed using MMseqs2 (17) using the cascaded clustering mode (mmseqs cluster, https://mmseqs.com/latest/userguide.pdf). Clustering was first performed on the proteins from the individual assemblies, and the cluster-representative proteins were subsequently clustered with the co-assembly proteins. The following parameters were used in the two MMseqs2 runs: -c 0.95; -- min-seq-id 0.95; --cov-mod 1; --clust-mod 2. This means proteins displaying ≥95% amino acid identity were clustered. Strains belonging to the same prokaryotic species generally display >95% average amino acid identity (18). As recommended in the MMseq2 user guide, -cov-mod 1 was used, since it allows clustering of fragmented proteins (as often occurs in metagenomic datasets). With --cov-mode 1 only sequences are clustered that have a sequence length overlap greater than the percentage specified by -c (i.e. 95% with -c 0.95) of the target sequence. In MMseqs2, the query is seen as the representative sequence, and the target is a member sequence. To lower the risk for fragmented proteins becoming cluster-representative sequences, -cluster-mode 2 was used, again following the recommendations of the MMseq2 user guide. It sorts sequences by length and in each clustering step forms a cluster containing the longest sequence and the sequences that it matches.

### Read mapping and counting

Random subsets of 10,000 non-normalized forward reads per sample were created using seqtk v.1.2-r101-dirty (https://github.com/lh3/seqtk), with seed 100 (-s 100). These reads (12.4 million in total) were mapped to the representative gene sequences from either the individual, co-, or mix-assembly using Bowtie2 v.2.3.4.3 (19), with the parameter “--local”. The resulting SAM files were converted to BAM with Samtools v.1.9 (20). The htseq-count script from HTSeq (21) v.0.11.2 was used to obtain raw counts per gene, with the parameters “-f bam -r pos -t CDS -i ID -s no -a 0”. For the counting, GFF input files were used, created using the script create_gff.py available at https://github.com/EnvGen/toolbox/tree/master/scripts. In order to estimate read depth coverage of the genes in the total metagenome, we multiplied the counts per gene by the average read-pair length divided by the length of the gene, and multiplied this number with the total number of read-pairs in the whole dataset divided by the total number of randomly sampled forward reads. This is a rough estimation of the coverage of each gene in the total metagenome, however after normalisation with BBnorm, high coverage genes will get a lower coverage.

### Functional annotations

Functional annotation of proteins were conducted using EggNOG (22), Pfam (23), and dbCAN (24). Annotations against Pfam v.31.0 and dbCAN v.5.0 were conducted with hmmsearch and hmmscan (25), respectively, in HMMER v.3.2.1, selecting hits with E-value < 0.001. Annotations against EggNOG v.4.5.1 were performed using eggNOG-mapper v.1.0.3 (26), using Accelerated Profile HMM Searches (27), following the recommendation for setting up large annotation jobs.

### Taxonomic affiliation

MMseqs2 (v13.45111) taxonomy (28), with parameters “--orf-filter 0 -- tax-lineage 1”, was used to assign taxonomic labels to contigs from which representative genes were predicted. MMseqs2 taxonomy uses an approximate 2bLCA (Lowest Common Ancestor, LCA) approach. GTDB (29, 30) v.202 was used as a reference database for Bacteria and Archaea and Uniprot90 (31) (downloaded on June 4th, 2021) for Eukaryota and Viruses.

### RNA gene screening

Barrnap v.0.9 (32), using default parameters, was used to identify potential rRNA genes, and identification of rRNA and other potential RNA genes in the mix-assembly gene set was conducted using the Rfam v.14.6 (33) database, with hmmsearch (25), in HMMER v.3.3.2, with flag “--cut_ga”. The union of genes identified as rRNA by Barnap and Rfam/hmmsearch were removed from the final gene set.

### Data availability

The shotgun reads and individual sample assemblies have been published earlier (6, 7, 12). The co-assembly contigs and the mix-assembly gene set (BAGS) together with annotations are available at the SciLifeLab Data Repository powered by Figshare, https://doi.org/10.17044/scilifelab.16677252. The contigs for the individual assemblies were published earlier (7) and are available at ENA hosted by EMBL-EBI under the study accession number PRJEB34883. When using the BAGS gene set in your work, please cite Alneberg et al. (2020)(7) in addition to this study.

## RESULTS

We used a set of 124 metagenome samples from the Baltic Sea ((6, 7, 12); Fig. S1) to evaluate three assembly approaches for generating a non-redundant gene catalogue: co-assembly on all samples (‘co-assembly’), assembly on individual samples (‘individual-assembly’), and a combination of the previous two (‘mix-assembly’). For the co-assembly, due to the complexity of the dataset, direct co-assembly of all reads was not possible, even on a server with 1 TB of memory. Therefore, the reads were first normalised such that reads stemming from highly abundant genomes (with high-frequency *k*-mers) were down-sampled (to a depth of 70x coverage), and those presumably derived from errors (with a depth below 2x) were removed. This reduced the total number of read-pairs from 5.4 to 2.9 billion.

Since the contigs of the co-assembly are derived from reads from all samples, it will result in a non-redundant set of genes. In contrast, genes from the individually assembled samples may overlap between samples. To reduce this redundancy, clustering was conducted on the encoded proteins (17). We used a cutoff of 95% amino acid identity, conforming to that strains belonging to the same species typically display more than 95% average amino acid identity (18). This reduced the number of individual-assembly genes from 134 to 50 million. Likewise, clustering was conducted on the co-assembly proteins together with the non-redundant set of individual-assembly proteins, to generate the mix-assembly gene set.

The mix-assembly approach resulted in the largest number of non-redundant genes (67 M), followed by individual assembly (50 M) and co-assembly (45 M; Table 1). Mix-assembly also had the largest number of genes predicted to be complete (12 M) followed closely by co-assembly (11 M), but twice as many as individual assembly (6 M; Table 1).

**Table 1.** Representative gene characterisation of different assembly approaches.

| **Assembly approach** | **Total bps** | **Number of genes** | **Num. of genes $\geq 100$ bp** | **Num. of Complete genes** | **Num. of Partial genes** | **Num. of Incomplete Genes** |
| --- | --- | --- | --- | --- | --- | --- |
| *Individual* | 18,770,879,205 | 50,045,582 | 45,859,319 | 6,258,868 | 27,073,554 | 16,713,160 |
| *Co* | 20,347,887,912 | 45,455,222 | 42,278,556 | 11,443,584 | 23,815,733 | 10,195,905 |
| *Mix* | 27,043,772,505 | 67,583,055 | 61,576,531 | 12,690,647 | 37,345,617 | 17,546,791 |

The gene size distributions were fairly similar for the three approaches (Fig. 1), with peaks in the distributions between 300 and 350 bp. Co-assembly had the largest median gene length (336 bp), although mix-assembly had the largest number of genes along the full range of gene sizes (Fig. 2).

**FIG 1.**
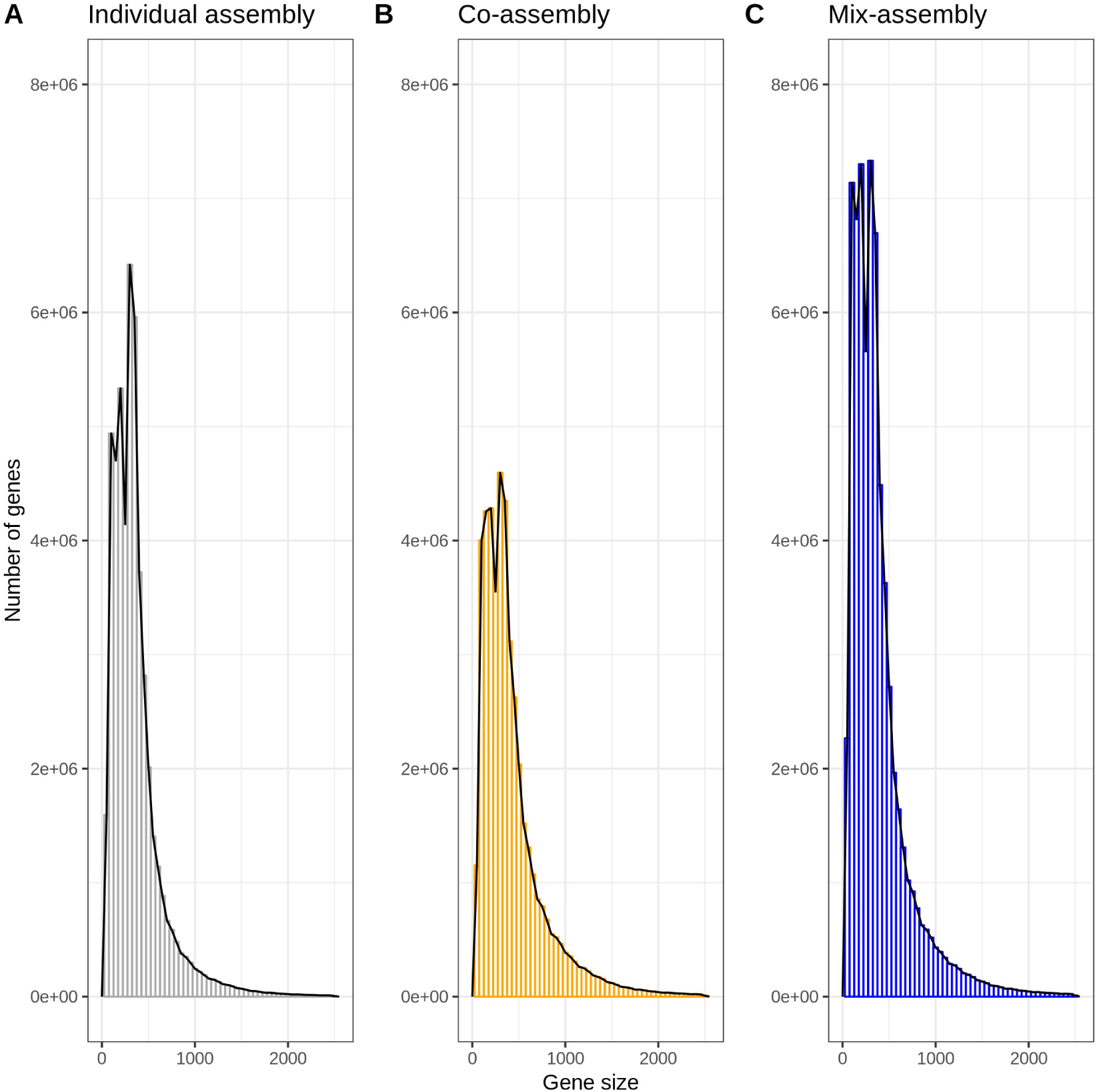
Gene size distributions of the three assembly approaches. (a) Co-assembly. (b) Individual-assembly. (c) Mix-assembly. Only genes ≤1500 bp are included in the histograms.

**FIG 2.**
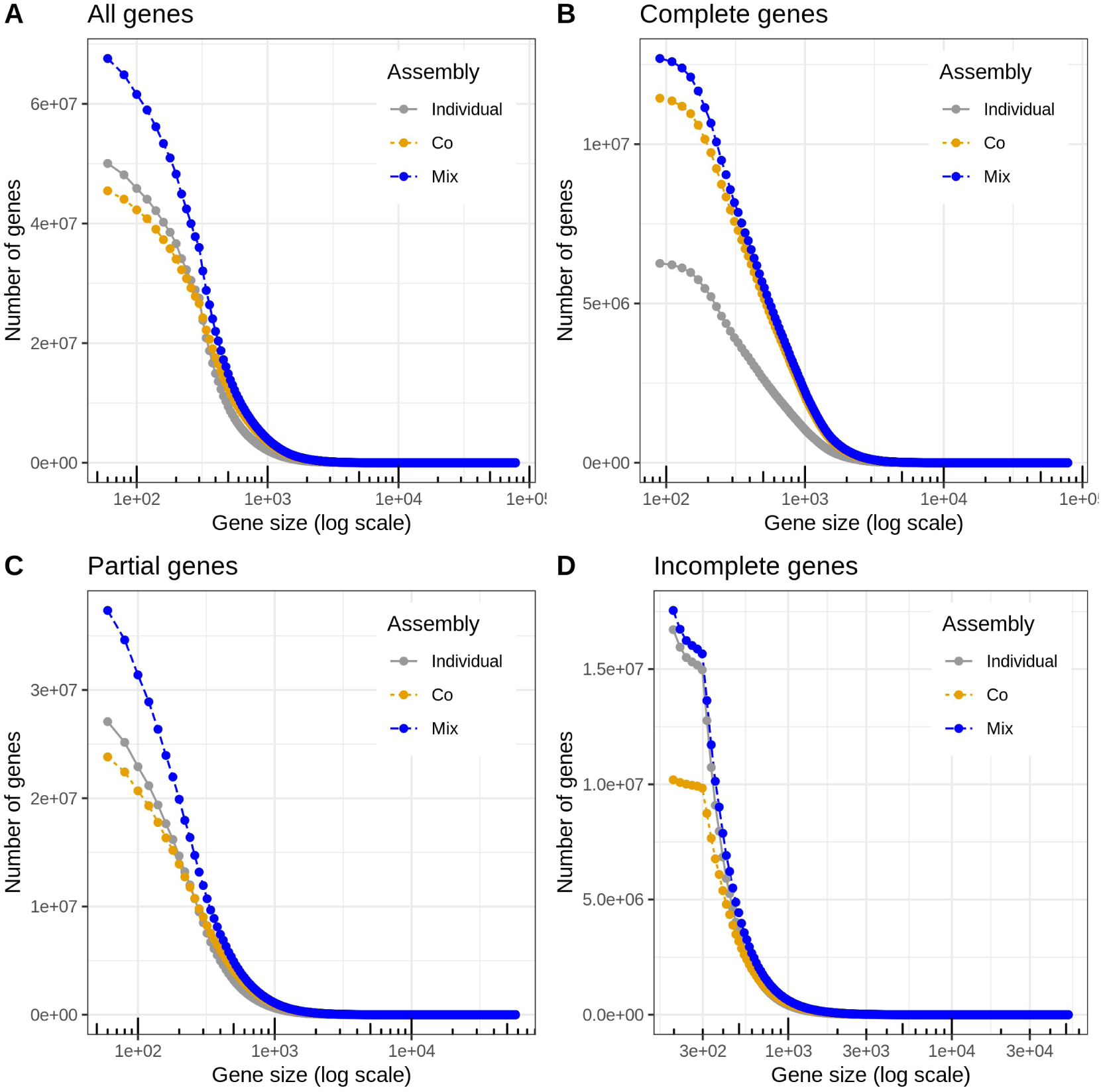
Cumulative distribution of gene sizes for the three assembly approaches. (a) All genes. (b) Complete genes. (c) Partial genes. (d) Incomplete genes.

Annotating the proteins against Pfam (23) gave the largest number of annotated genes for mix-assembly (15 M) followed by co-assembly (13 M) and individual-assembly (12 M), despite that co-assembly had a higher proportion of genes with annotation (29.4%) compared to the other two (23.0% for mix-assembly, 23.8% for individual assembly; Table 2).

**Table 2.**
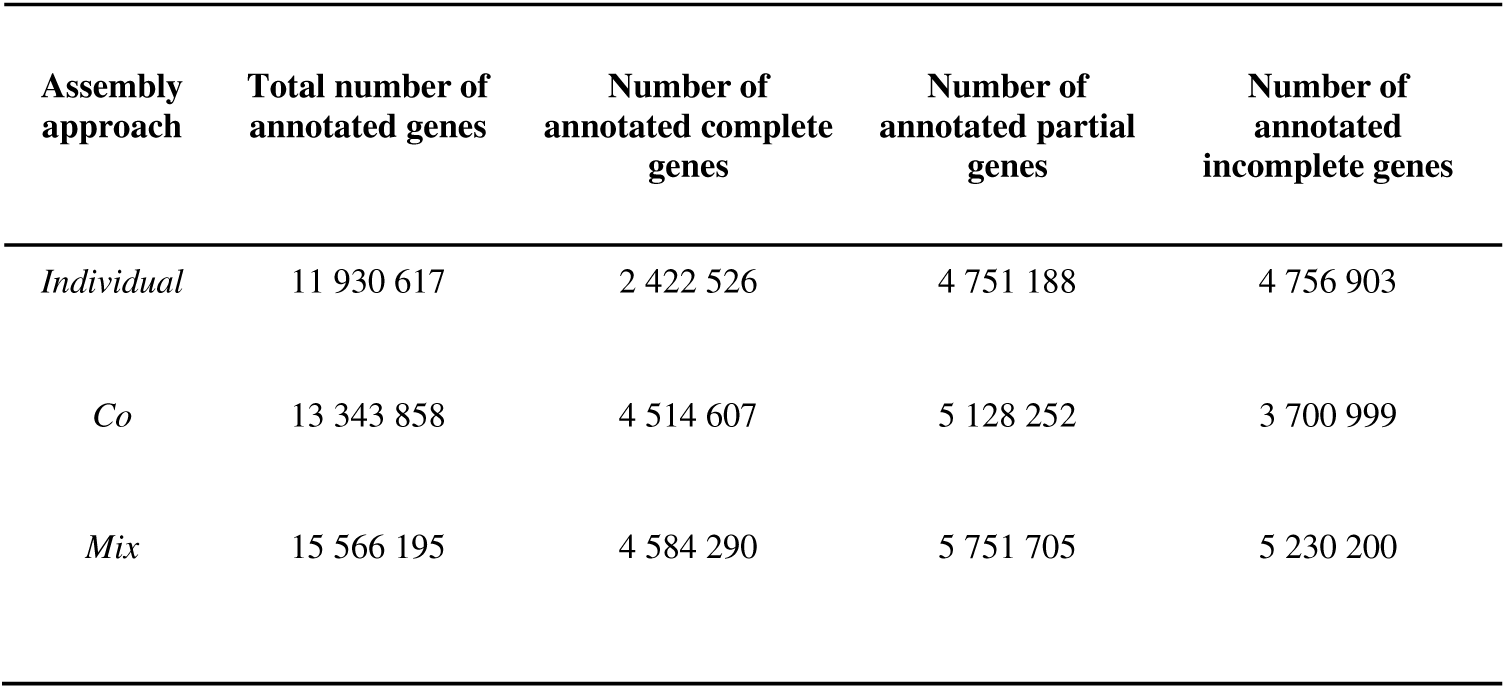
Pfam annotation of representative proteins from different assembly approaches.

Since biome-specific gene catalogues are often used as reference sequences for mapping of shotgun reads from metagenomes or transcriptomes, we further evaluated the gene sets by mapping reads from the metagenome samples to them. The average mapping rates for the 124 samples were 83.9, 84.7, and 87.7% for individual-, co- and mix-assembly, respectively, with numbers ranging from 47.5, 49.2 and 53.2% to 96.2, 96.1 and 97.3% for individual-, co- and mix-assembly. The mix-assembly read-mapping rate was significantly higher than the individual- (Wilcoxon rank-sum test, *P* < 10^−5^) and co-assembly (*P* < 10^−4^) rates (Fig. 3a). Fig. 4 presents the cumulative mapping rate by gene size, showing the proportion of reads mapping at different gene length cut-offs. For all three assembly strategies, the highest fraction of reads mapping corresponds to complete genes, followed by partial genes. Of the three, mix-assembly had the highest fraction of mapping reads mapping to complete genes (42.6%), and the lowest to partial (32.0%) and incomplete (13.1%) genes. Mix-assembly also had the highest proportion of reads mapping to genes with a Pfam annotation (56.9%, p.adj.value = 0.052 - Wilcoxon rank-sum test - p-value adjust method FDR), followed by co-assembly (54.0%) and individual-assembly (54.0%)(Fig. 3b).

**FIG 3.**
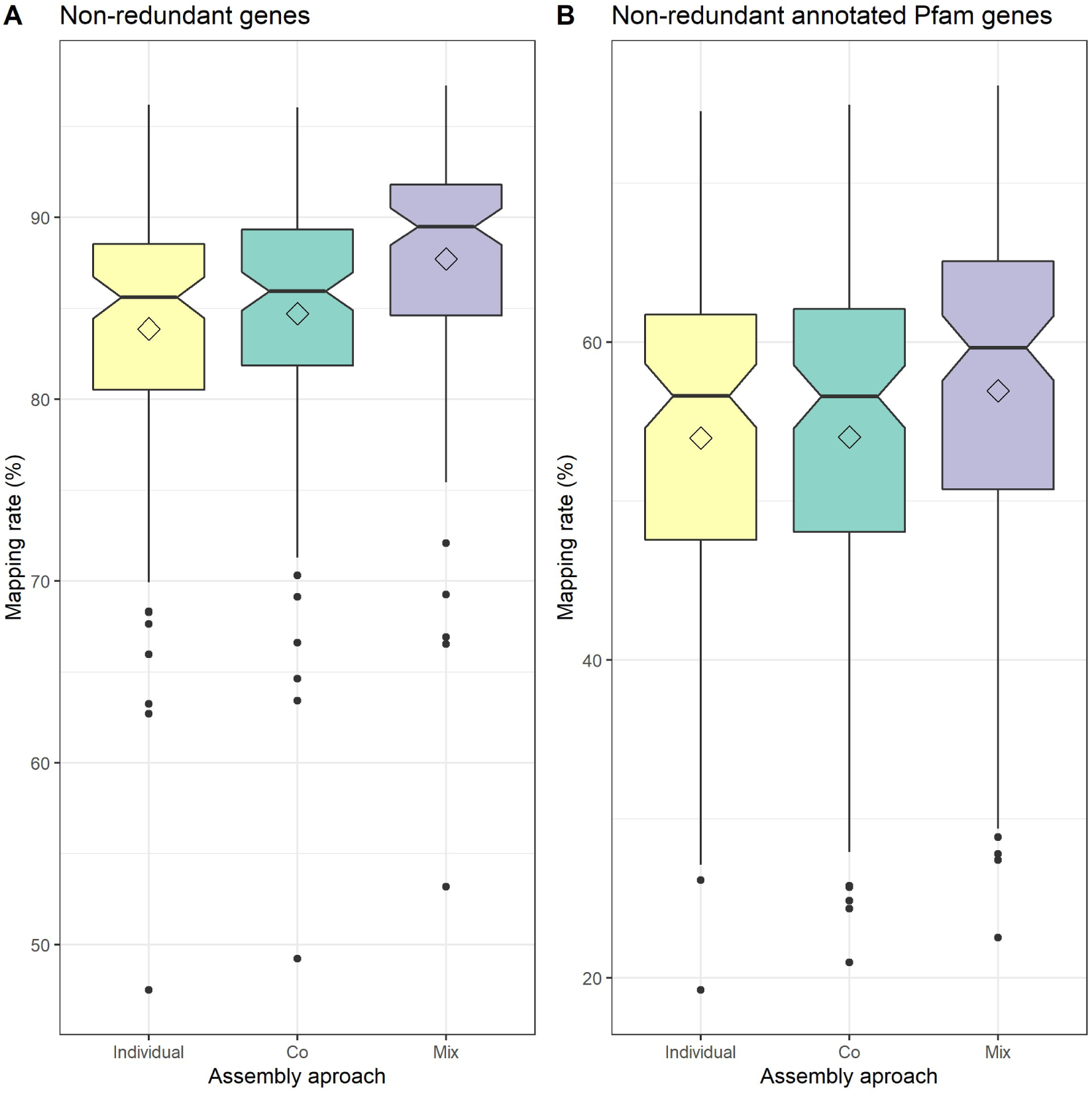
Read mapping rates to genes from the three assembly approaches. The boxplots show the distribution of mapping rate (% of reads) for the 124 samples, based on a random subset of 10,000 forward reads per sample. (a) For all genes. (b) For genes with Pfam annotation.

**FIG 4.**
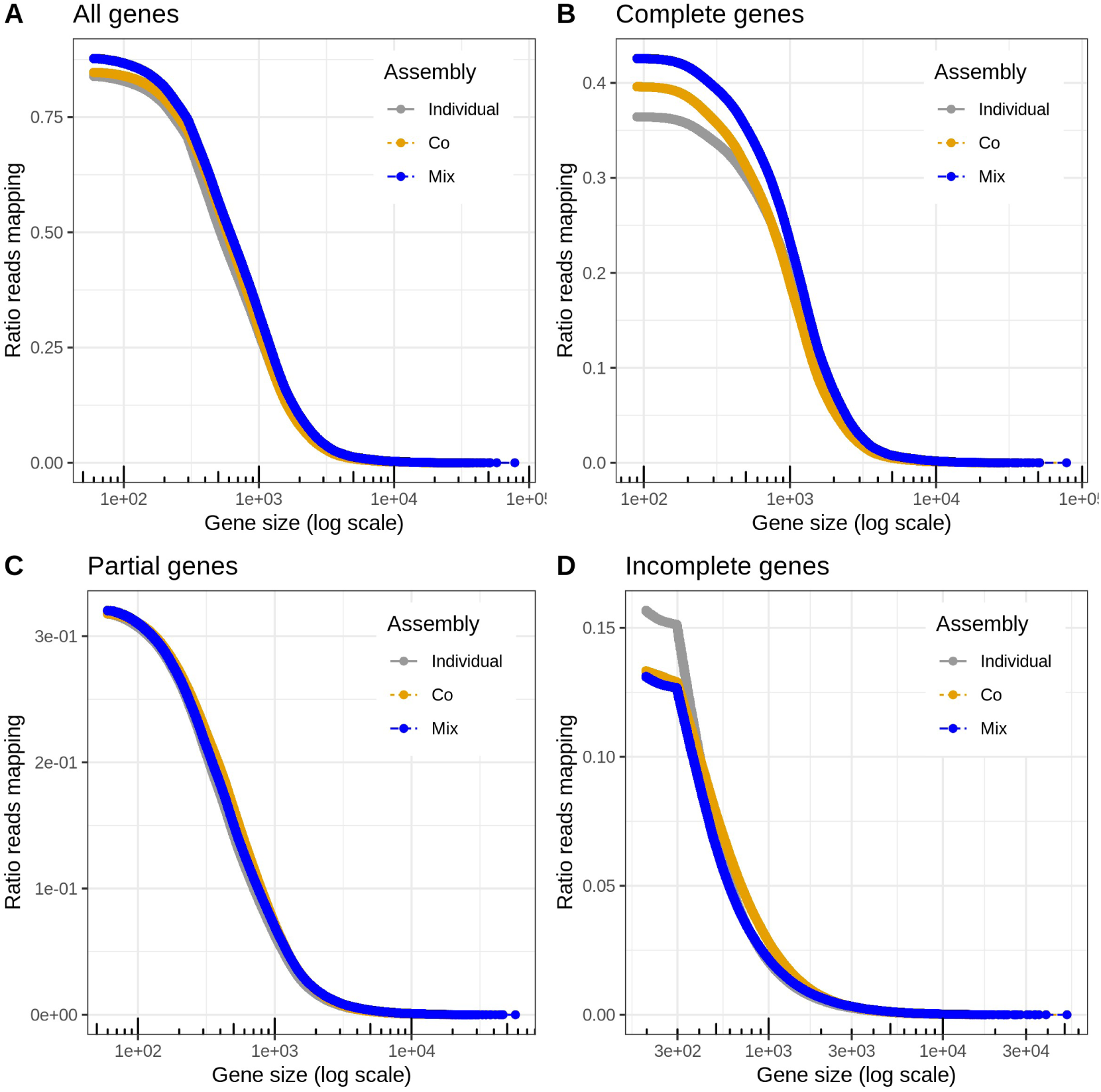
Read mapping rate as a function of gene length cut off. The plots show the ratio of reads mapping at different cut-offs on minimum gene length. (a) All genes. (b) Complete genes.(c) Partial genes. (d) Incomplete genes.

The contribution of genes from the individual- and co-assembly to the mix-assembly set of genes is shown in Fig 5. A majority (52%) of the mix-assembly genes originates from co-assembly genes (Fig. 5a), representing 67% of the complete and 50% and 45% of the partial and incomplete genes, respectively (data not shown). However, among the reads that map to the mix-assembly genes, a larger fraction of reads map to genes derived from the individual-assembly than to genes derived from the co-assembly (Fig. 5b). These seemingly conflicting results may reflect that mix-assembly genes derived from the individual-assembly tend to be of higher abundance in the microbial communities than those from the co-assembly. This was confirmed by grouping the mix-assembly genes in low, median and high coverage genes, where the majority of mapping reads mapped to genes derived from co-assembly for low coverage genes but to genes derived from individual-assembly for high coverage genes (Fig. 5c).

**FIG 5.**
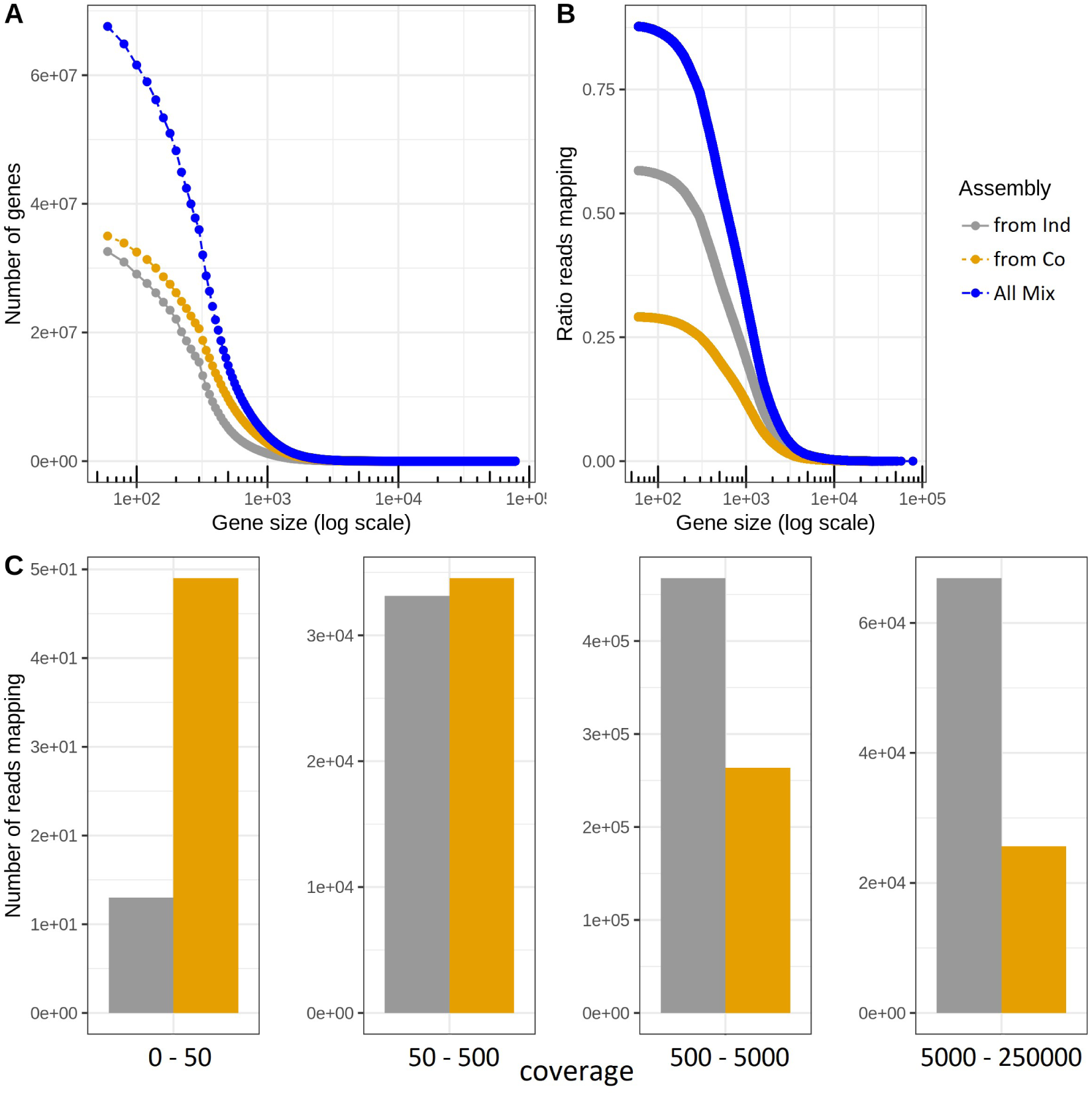
Contribution of genes from individual-assembly and co-assembly to the mix-assembly gene set. (a) Cumulative distribution of gene sizes for the mix-assembly genes: for all (‘All Mix’) and for those derived from individual-assembly (‘from Ind’) and co-assembly (‘from Co’). (b) Read mapping rate as a function of gene size cut off.(c) Total number of reads mapping to mix-assembly genes derived from either individual-assembly or co-assembly, for four bins of genes binned by their estimated coverage in the total metagenome (see Methods): low (0 - 50 x), median (50 - 500 x), high (500 - 5,000 x) and very high (5,000 - 250,000 x) read depth coverage.

The mix-assembly gene set is significantly more extensive than the previously published Baltic Sea gene catalogue (BARM;(6)) and may serve as a valuable resource for brackish water research We compared the mix-assembly protein set with the Tara Ocean Microbial Reference Gene Catalog (OMG-RGC.v2 (34)). Of the 67.5 M representative mix-assembly proteins, only 1.4 M were >95% identical to Tara proteins, and vice versa, of the 46.7 M Tara proteins, 1.3 M were >95% identical to the representative mix-assembly proteins. Hence, the vast majority of the mix-assembly gene sequences are distinct from Tara genes. To increase the usefulness of the mix-assembly gene set, we removed genes potentially encoding ribosomal RNA and thus falsely predicted as protein-coding (n=16,804), and conducted taxonomic and functional annotation on the remaining genes. A subset of the genes (n=70,223) was predicted to include encodings of other structural RNAs (in Rfam (33)), but we decided to keep these since they may also encode important protein-coding regions. The resulting gene set, that we call BAltic Gene Set (BAGS.v1), encompasses 67,566,251 genes, of which 31.0 M have a taxonomic affiliation (Fig. S2) and 23.4 M have at least one type of functional annotation: 15.5 M with PFAM, 21.5 M with EggNOG (22), 1.5 M with dbCAN (24) annotation (Table 3). Twentyseven percent of the BAGS.v1 genes were predicted to be of eukaryotic origin. It should however be noted that the gene predictions were conducted with a gene caller for prokaryotic genes (Prodigal) and that a fraction of the eukaryotic genes has likely been imperfectly predicted.

**Table 3.** Number of mix-assembly representative genes annotated against several databases.

| **Gene completeness** | **dbCAN** | **EggNOG** | **Pfam** |
| --- | --- | --- | --- |
| *complete* | 420 422 | 5 354 169 | 4 582 506 |
| *partial* | 562 445 | 8 374 034 | 5 751 622 |
| *Incomplete* | 603 580 | 7 865 395 | 5 230 173 |
| **TOTAL** | **1 586 447** | **21 593 598** | **15 564 301** |

## DISCUSSION

Metagenome assembly is commonly carried out either by individually assembling reads from each sample (35) or by co-assembling reads from all the samples of a dataset (2, 6). Here, the performance of these assembly approaches was compared. Although the number of genes was lower for the co-assembly, the total length (in number of base pairs) was higher than for the individual assembly. The two gene sets reported a similar mapping rate, although the co-assembly set had a higher number of genes predicted to be complete and a lower number of partial and incomplete genes than the individual-assembly set. In this study, we also proposed a new approach for assembly, aiming to combine the advantages of the individual- and co-assembly approaches, referred to as mix-assembly. The mix-assembly strategy resulted in significantly (35 and 48%) more genes than the other approaches and also in the largest number of complete genes. It further gave the highest mapping rates and the greatest number of genes with a Pfam annotation. The reason why not only the number of genes, but also the number of complete genes increased compared to the other approaches, is likely because in the protein clustering process the longest proteins were selected to form cluster seeds. Thus, if for example an incomplete or partial protein from the co-assembly set forms a cluster with a complete protein from the individual-assembly, the complete protein will likely represent this cluster in the mix-assembly, since it is longer. Thereby, the clustering step that combines the two gene sets enriches for complete proteins. However, it may also to some extent enrich for artificially long proteins that may stem from sequencing or gene calling errors.

Analysing the contribution of individual- and co-assembly genes in the set of mix-assembly genes showed that genes with relatively low coverage (low number of mapping reads) in the samples were mainly stemming from the co-assembly. This likely reflects that co-assembly sometimes is able to recover genes that display too low coverage to be assembled from individual samples. On the other hand, genes with relatively high coverage were mostly originating from the individual-assembly, which may be caused by the co-assembly sometimes breaking in such genes due to strain variation. If strain variation for such a gene is less pronounced in at least one of the individual samples, a longer fraction of the gene could be recovered in the individual-assembly.

The 67 million genes of the mix-assembly are based on 124 metagenome samples that span the salinity and oxygen gradients of the Baltic Sea and also capture seasonal dynamics at two locations (7). This dataset (BAGS.v1) is a 10-fold expansion compared to our previous gene set (6) and has the potential to serve as an important resource for exploring gene functions and serve as a backbone for mapping of meta-omics data from brackish environments. Consistent with our earlier study showing that the prokaryotes of the Baltic Sea are closely related to but genetically distinct from freshwater and marine relatives (35), only a small fraction of the mix-assembly genes displayed >95% amino acid similarity to genes of the Tara Ocean gene catalogue. This implies that the Tara Ocean catalogue is not suitable for mapping of meta-omics data from the Baltic Sea and emphasizes the need for a brackish water microbiome reference gene catalogue. The gene catalog BAGS.v1, including gene and protein sequences, and taxonomic and functional annotations, is publicly available at the SciLifeLab Data Repository, https://doi.org/10.17044/scilifelab.16677252.

## ACKNOWLEDGEMENTS

This work is part of the Swedish Biodiversity Data Infrastructure (SBDI; https://biodiversitydata.se), funded by its partner organizations and the Swedish Research Council VR through Grant No 2019-00242. Computations were performed on resources provided by the Swedish National Infrastructure for Computing (SNIC) through the Uppsala Multidisciplinary Center for Advanced Computational Science (UPPMAX).

## FIGURE LEGENDS

**FIG S1** Map with sampling locations. The marker colour shows the salinity of the water sample and its size, the sampling depth. The contour lines indicate depth with 50 m intervals.

**FIG S2** BAG interactive taxonomic affiliation figure. Available at the SciLifeLab Data Repository, https://doi.org/10.17044/scilifelab.16677252

## TABLE FOOTNOTES

**TABLE 1** Assembly and gene statistics of the different assembly approaches.

**TABLE 2** Statistics on Pfam annotations for the different assembly approaches.

**TABLE 3** Statistics on mix-assembly proteins annotated against different databases.

**TABLE S1** Sample retrieval and sequencing description (further sample description in references).

